# Paired scTCR-seq as a Complementary Reference for RNA-Based Doublet Detection

**DOI:** 10.64898/2026.11.27.769697

**Authors:** Yu-Ting Weng, Shih-Hsiang Lo, Yi-Hui Huang, Sheng-Kai Lai, I-Hsuan Chiu, Pei-Lung Chen, Chien-Yu Chen

**Author notes:** Corresponding authors: Pei-Lung Chen and Chien-Yu Chen. These authors contributed equally (Yu-Ting Weng, Shih-Hsiang Lo).

## Abstract

Computational doublet detection is a routine step in single-cell RNA sequencing (scRNA-seq), but real datasets lack the ground truth needed to check it. Paired single-cell T-cell receptor sequencing (scTCR-seq) offers a signal that is independent of the features used by RNA-based algorithms: a barcode carrying three or more productive chains of either TRA or TRB is unlikely to contain a single T cell. We compared six RNA-based doublet-detection methods with this pipeline-derived TCR potential-multiplet flag in 96,003 cells from 44 Parse Evercode peripheral blood samples, of which 70,959 had detectable TCR sequences and 6,208 were flagged. Flag prevalence rose with RNA-tool agreement, from 2.6% among cells called by no method to 50.2% among cells called by four, before declining at five and six; the association was attenuated but persisted after adjustment for within-sample RNA depth. Methods differed several-fold in call rate, and requiring four rather than three positive methods increased precision against the flag by 4.3 percentage points but reduced recall by 22.7 points. Discordance occurred in both directions. The 1,453 flagged cells called by no RNA method had RNA content close to that of non-flagged TCR-positive cells called by no method (median UMI count, 6,549 versus 6,227), so their multichain signal was not accompanied by the RNA excess typical of conventional multiplets. Conversely, of 208 non-flagged cells called by all six methods, 32.2% carried B-cell, myeloid, or platelet signal, compared with 10.5% of 54,914 cells called by none (sample-stratified odds ratio, 4.08; 95% confidence interval, 3.03–5.50), consistent with T–non-T mixtures that a chain-count rule cannot detect; most of the remainder carried about twice the RNA content of reference cells. Paired scTCR-seq therefore provides a complementary reference that supports RNA-based consensus, while each modality reveals events that the other misses. In immune-profiling data, combining both and retaining the consensus count as graded evidence offers a more complete approach to doublet detection.

## 1 Introduction

Doublets arise when two or more cells receive the same single-cell barcode. Their combined transcriptomes can resemble intermediate or rare cell states, distort cell-type proportions, and bias clustering, annotation, differential expression, and trajectory analyses. Because the true identities of captured cells are generally unobserved, computational doublet detection has become a standard component of single-cell RNA sequencing (scRNA-seq) workflows.

Existing methods use different signals, including simulated doublets, neighborhood structure, improbable gene-pair co-expression, and latent representations (Wolock et al., 2019; McGinnis et al., 2019; Bais and Kostka, 2020; Bernstein et al., 2020; Germain et al., 2021; Gayoso and Shor, 2025). Their predictions can therefore differ substantially even when applied to the same count matrix. Prior benchmarking has also shown that no method dominates across all datasets and evaluation criteria (Xi and Li, 2021). Experimental labels used for benchmarking are themselves incomplete: genotype demultiplexing, species mixing, cell hashing, and lipid-tag multiplexing identify different subsets of multiplets and may fail to label homotypic events.

Paired single-cell T-cell receptor sequencing (scTCR-seq) offers an orthogonal signal in T-cell-rich samples. Because each TCR locus has two alleles, a single T cell is expected to express at most two productive alpha (TRA) or beta (TRB) chains, and because TRB undergoes stricter allelic exclusion than TRA, additional TRB chains are less readily explained by a single T cell. A barcode carrying three or more productive chains of either type is therefore unlikely to contain a single T cell and can serve as a potential-multiplet label that does not depend on the RNA-expression features used by doublet-detection algorithms.

This label has its own blind spots. It is informative only for cells with detectable TCR sequences, and it is expected to be most sensitive to T–T multiplets: a mixture of a T cell with a B cell or myeloid cell, or of two conventional T cells, may carry too few chains to be flagged. Technical events that add apparent chains could also flag a single cell (Fig. 1). TCR multiplicity should consequently be interpreted as a complementary cross-modal reference rather than complete doublet ground truth.

**Figure 1:**
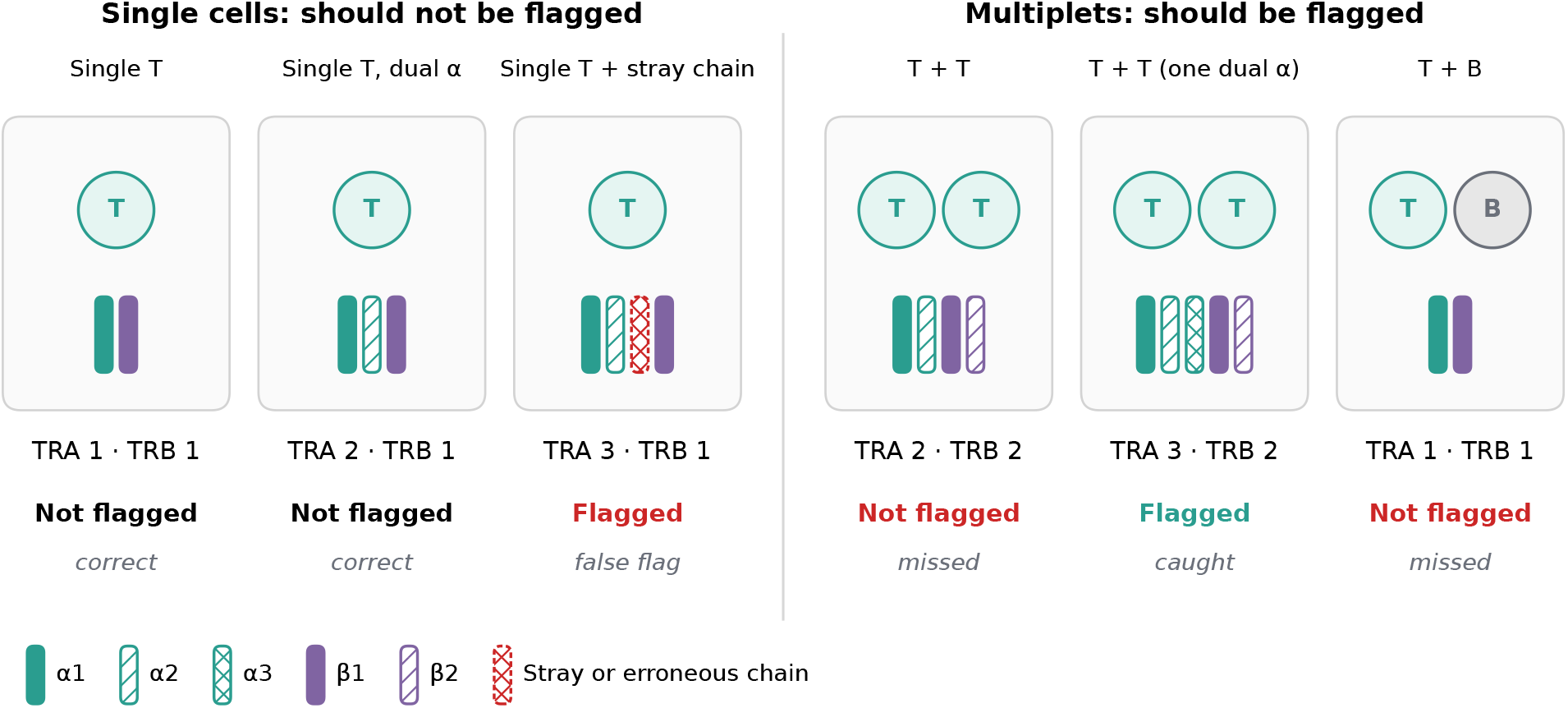
Expected behavior of the three-chain TCR potential-multiplet rule. A barcode is flagged when it carries three or more productive TRA chains or three or more productive TRB chains. Single T cells, including dual-*α* T cells, should not be flagged, but an additional stray or erroneous chain can produce a false flag. Among multiplets, two conventional T cells contribute only two TRA and two TRB chains and are missed, whereas a T cell mixed with a non-T cell contributes no additional chains and is also missed. Chain configurations are illustrative; each fill pattern denotes a distinct chain sequence.

Immune-repertoire analysis frameworks such as scirpy similarly classify multichain receptor configurations as potential doublets (Sturm et al., 2020), and immune-receptor and surface-protein evidence has been used to label training doublets for a transcriptome-based classifier (Sun et al., 2021). Rather than using TCR evidence as a filter or as training labels, we treat it as a complementary reference and systematically examine where multichain TCR evidence and six RNA-based doublet-detection methods agree and where they disagree.

Here, we evaluated scDblFinder, cxds, DoubletFinder, Scrublet, DoubletDetection, and Solo in 44 Parse Evercode samples with paired scTCR-seq. We quantified method-specific concordance with TCR potential-multiplet flags, examined multi-tool consensus thresholds with supporting evidence from 16 public benchmark datasets, and characterized discordance in both directions. We asked three questions: whether the TCR flag became more frequent as more RNA-based methods agreed; what characterized TCR-flagged cells that no RNA-based method called; and whether unanimous RNA calls that lacked a TCR flag carried transcriptional signals of non-T lineages.

## 2 Methods

### 2.1 Study dataset and preprocessing

We analyzed a Parse Evercode single-cell RNA sequencing dataset comprising 96,003 quality-controlled cells from 44 samples and 29 participants. The dataset contained raw gene-level unique molecular identifier (UMI) counts, sample and participant identifiers, quality-control metrics, T-cell receptor (TCR) annotations, and the per-cell calls of six computational doublet-detection methods. Preprocessing, quality control, integration of TCR records, execution of the six methods (Section 2.2), and the public benchmark (Section 2.5) were performed by Y.-T.W. as part of her master’s thesis research; all subsequent analyses reported here were performed for the present study. All cells included in the analysis had passed the quality-control workflow defined in the original preprocessing pipeline, but computationally predicted doublets had not yet been removed.

The analysis began with 101,689 cells generated using the Parse Biosciences Evercode Whole Transcriptome v3 platform, which uses split-pool combinatorial barcoding (Rosenberg et al., 2018), and processed with the Parse Biosciences split-pipe pipeline (version 1.3.1 for RNA data). Cells with a sample-specific number of detected genes or total UMI count below the median minus three median absolute deviations were removed, as were cells with mitochondrial transcripts exceeding 5% of total counts. Upper outliers in genes or UMIs were recorded but were not removed at this stage, thereby avoiding the use of RNA burden as a pre-emptive doublet filter. DecontX contamination estimates (Yang et al., 2020) were retained for quality assessment but were not used as exclusion criteria. The resulting dataset contained 96,003 cells; the number per sample ranged from 1,022 to 5,642 (median, 1,964; mean, 2,182). Raw count matrices were retained for doublet detection and marker-expression analyses.

Peripheral blood mononuclear cell samples were provided by the laboratory of Pei-Lung Chen at the College of Medicine, National Taiwan University. The cohort comprised patients with Graves’ disease (GD), patients with antithyroid drug-induced agranulocytosis (TiA), and healthy controls. Patients with GD (*n* = 9) and TiA (*n* = 6) each contributed paired samples collected during the acute and recovery phases, and healthy controls (*n* = 14) contributed one sample each, yielding 44 samples from 29 participants. Further details of the cohort are described in Huang et al. (2025).

### 2.2 Computational doublet detection

Six computational doublet-detection methods were evaluated: scDblFinder, cxds, DoubletFinder, Scrublet, DoubletDetection, and Solo. Each method was applied independently to each sample to avoid confounding caused by differences in library size, cell loading, and cell-state composition across samples.

For methods requiring an expected doublet rate, a nominal rate of 3% was specified. This value was used as an algorithmic input and was not interpreted as the true biological doublet prevalence. cxds did not use this fixed rate; its doublet number was estimated internally with estNdbl=TRUE. A fixed random seed of 666 was used for stochastic procedures, including simulated-doublet generation, sampling, and model training.

scDblFinder (version 1.16.0) was run in pure-random artificial-doublet mode with propRandom=1 and an expected doublet rate of 0.03; its internally optimized threshold produced the binary classification (Germain et al., 2021). cxds, implemented in scds version 1.18.0, used the 500 highest-ranked genes and binarized expression at counts greater than zero; the package-generated cxds_call was used as its binary classification (Bais and Kostka, 2020). DoubletFinder (version 2.0.6) was run after Seurat normalization (Stuart et al., 2019), selection of 2,000 variable genes, scaling, and principal-component analysis using the first 30 components (McGinnis et al., 2019). It used *pN* = 0.25; the neighborhood-size parameter *pK* was selected separately for each sample by maximizing the mean–variance normalized bimodality coefficient. The expected number of doublets was adjusted for the estimated homotypic-doublet proportion before producing the binary classification.

Scrublet (version 0.2.3) was run using an expected doublet rate of 0.03, 30 principal components, and a simulated-doublet ratio of 2. Sample-specific thresholds derived from the simulated score distributions were used whenever the automatic thresholding procedure converged; a score threshold of 0.25 was used as a fallback (Wolock et al., 2019). DoubletDetection (version 4.3.0.post1) (Gayoso and Shor, 2025) was run for 50 iterations using Louvain clustering, a hypergeometric *P* -value threshold of 10*^−^*^6^, and a voting threshold of 0.5. Ambiguous predictions were not counted as doublets. Convergence logs indicated that 11 of 44 sample-level runs did not meet the stopping criterion at 50 iterations. Solo, implemented in scvi-tools version 1.3.3 (Gayoso et al., 2022), was trained using an scVI representation with two neural-network layers, 30 latent dimensions, and at most 400 epochs (Bernstein et al., 2020). Genes detected in fewer than three cells were excluded, and 2,000 highly variable genes were retained. A cell was classified as a doublet when its predicted doublet probability exceeded its singlet probability.

### 2.3 Consensus doublet classification

Because no single method performs best across datasets (Xi and Li, 2021), we summarized the six methods as a consensus score. For each cell, we calculated the number of methods that classified it as a doublet, producing a consensus score ranging from zero to six. A score of zero indicated that no method classified the cell as a doublet, whereas a score of six indicated unanimous classification by all six methods.

Consensus classifications were defined as at least *k* positive calls, where *k* ranged from one to six. Informed by benchmarking against experimentally annotated public datasets, agreement by at least four methods was used as the reference operating threshold. Cells receiving all six positive calls were additionally examined as a unanimous RNA-call group.

The homotypic-adjusted DoubletFinder classification was used in all consensus calculations.

### 2.4 TCR potential-multiplet flag as a cross-modal reference

The Parse-derived `tcr_is_multiplet` field was used as a cross-modal reference for evaluating RNA-based calls. According to the Parse Biosciences pipeline documentation, the flag denotes barcodes with three or more productive TRA chains or three or more productive TRB chains; because each TCR locus has two alleles, a threshold of three avoids flagging single T cells that express two productive chains of one type (Parse Biosciences, 2023). Throughout the manuscript, cells carrying this flag are termed *TCR potential multiplets*, rather than true doublets. The flag is restricted to cells with informative TCR data and may capture TCR-specific technical events, such as ambient or misassigned TCR transcripts or sequence variants of one chain called as distinct chains, in addition to physical cell multiplets. Conversely, a multiplet composed of two conventional T cells can contain two TRA and two TRB chains and may not satisfy a rule based on three or more chains of either type. It therefore does not constitute complete or definitive doublet ground truth.

TCR libraries were generated with the Parse Evercode TCR assay from the same barcoded cells. TCR records were matched to RNA cells within each sample using the three combinatorial barcode indices and the sublibrary identifier, and a cell was considered to have detectable TCR information when a matching TCR record existed. Among the 96,003 cells, 70,959 had detectable TCR information and 6,208 met the TCR-multiplet definition. Cells whose barcodes did not appear in the TCR report were not treated as TCR singlets.

At each RNA-tool agreement level, TCR potential-multiplet prevalence was calculated only among cells with detectable TCR information. Binomial 95% confidence intervals were calculated using the Wilson method.

For each computational method and consensus threshold, discrimination of TCR multiplets was summarized using precision, recall, F1 score, and the area under the precision–recall curve (AUPRC), as applicable. AUPRC used each method’s continuous score: the scDblFinder score, the cxds score, the DoubletFinder proportion of artificial nearest neighbors (pANN), the Scrublet doublet score, the DoubletDetection doublet score, and the Solo doublet probability; the same score types were used in the public benchmark. In the Parse cohort, AUPRC was computed as average precision using scikit-learn. Mean sample-level areas under the precision–recall curve and pooled precision, recall, and F1 scores for the three- and four-tool consensus thresholds were assigned 95% percentile confidence intervals using 2,000 sample-cluster bootstrap replicates. Paired bootstrap differences between the two consensus thresholds were also calculated.

### 2.5 Public benchmark datasets

The six methods and their consensus classifications were additionally evaluated using 16 experimentally annotated benchmark datasets curated by Xi and Li (2021). Together, these datasets contained 168,239 cells, including 25,780 labeled doublets, and represented species mixing, cell hashing, genotype demultiplexing, and lipid-tag multiplexing. In this secondary benchmark, stochastic tools were run with seeds 42 and 1234 and their continuous scores were averaged. Performance was compared using metrics robust to class imbalance, with particular emphasis on Matthews correlation coefficient and precision–recall performance. In this benchmark, AUPRC was computed by integration of the precision–recall curve using the R package PRROC; because this estimator differs from average precision, AUPRC values are compared between the two settings only as within-setting tool rankings. Dataset sources and accessions are listed by Xi and Li (2021), and complete benchmark parameters are available from the corresponding authors upon request.

Binary calls in the public benchmark were generated using a dataset-specific threshold selected to maximize F1 score against the experimental labels for each method. The resulting consensus analysis is therefore an oracle-threshold benchmark rather than a direct simulation of unsupervised deployment. Under this framework, the threshold requiring at least four positive methods achieved the highest mean Matthews correlation coefficient. In the Parse cohort, package-derived or prespecified operating thresholds were used instead. The transportability of the four-tool threshold across these settings was treated as a limitation pending deployment-style or cross-validated sensitivity analysis.

### 2.6 RNA complexity across agreement levels

To assess whether shared dependence on RNA content could explain the association between RNA-tool agreement and the TCR flag, total UMI count and number of detected genes were converted to within-sample percentile ranks and grouped into quintiles. Flag prevalence was summarized by agreement level within each quintile, and the flag was modeled by logistic regression on the number of positive methods (as a numeric variable) with sample fixed effects, with and without quintile indicators. Odds ratios per additional positive method were assigned 95% percentile intervals from 2,000 sample-cluster bootstrap replicates.

TCR potential multiplets were grouped by the number of RNA-based methods classifying each cell as a doublet (zero through six). Total UMI count and number of detected genes were summarized by median and interquartile range. Associations between agreement level and RNA complexity were quantified using Spearman rank correlations. To reduce confounding by between-sample sequencing-depth differences, each complexity measure was additionally converted to a percentile rank within sample before correlation analysis. TCR potential multiplets receiving zero RNA-based calls were also compared with TCR-positive cells without the potential-multiplet flag using two-sided Mann–Whitney *U* tests and were described alongside TCR potential multiplets receiving at least one RNA-based call.

To distinguish a general association between RNA complexity and computational agreement from an association specific to the TCR flag, the within-sample percentile-rank correlations were calculated separately among TCR potential multiplets and TCR-positive cells without the flag. Differences between these correlations were quantified by cluster bootstrap resampling of samples and, separately, participants. Total UMI counts of unanimous RNA calls without high non-T-lineage signal (defined below) were compared with those of reference cells as a median ratio, with a 2,000-replicate sample-cluster bootstrap interval, and as a within-sample percentile rank among TCR-positive, non-flagged cells.

### 2.7 Chain-level audit of the TCR potential-multiplet flag

Primary and secondary TRA and TRB read counts, transcript counts, and chain annotations were examined in three prespecified groups: TCR-positive cells without the potential-multiplet flag, flagged cells receiving zero RNA-based calls, and flagged cells receiving at least one RNA-based call. For each cell, total TCR evidence was defined by summing available primary and secondary TRA and TRB read counts or transcript counts. Because only primary and secondary chains are reported for each locus, this sum may omit additional chains in flagged barcodes and is a truncated summary rather than a complete count of TCR molecules. Transcript counts, which collapse reads sharing a unique molecular identifier assigned before amplification, were used as the primary measure of TCR evidence because read counts additionally depend on targeted TCR enrichment and on the sequencing depth of the TCR library; read counts were summarized as a secondary measure. The prevalence of secondary TRA, secondary TRB, and both secondary chains was also summarized. Median differences and percentage-point differences were assigned 95% confidence intervals by 2,000-replicate cluster bootstrap, resampling samples and participants in separate analyses.

Because total TCR evidence sums chains that also contribute to the flag definition, these chain-level summaries were interpreted as descriptions of chain configuration rather than as independent validation of the flag. To assess whether TCR evidence scaled with transcriptome capture, within-sample Spearman correlations between total UMI count and total TCR reads or transcripts were computed among non-flagged TCR-positive cells. Because these associations were weak, chain-level comparisons are reported without adjustment for RNA complexity.

### 2.8 Exploratory lineage-marker analysis of unanimous RNA calls

To characterize cells unanimously classified as doublets despite lacking a TCR potential-multiplet flag, we performed a targeted marker-expression analysis. The target group consisted of TCR-detected cells that did not carry the TCR potential-multiplet flag but were classified as doublets by all six computational methods:

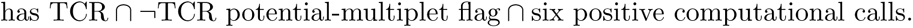

The reference group consisted of TCR-detected, non-TCR-multiplet cells that were not classified as doublets by any of the six methods. The target and reference groups contained 208 and 54,914 cells, respectively.

Raw UMI counts were library-size normalized to 10,000 counts per cell and log-transformed:

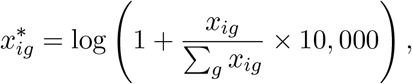

where *x_ig_* denotes the raw count for gene *g* in cell *i*. For each lineage, a lineage score was calculated as the mean normalized expression of its prespecified marker genes, a simplified form of gene-set scoring (Tirosh et al., 2016) without control-gene subtraction; thresholds were instead defined relative to reference cells within each sample, as described below.

The marker sets were:

- T cell: CD3D, CD3E, TRAC, CD2, CD247, and LCK;
- B cell: MS4A1, CD79A, CD79B, CD19, CD22, and CD37;
- natural killer cell: NKG7, GNLY, KLRD1, PRF1, and CTSW;
- myeloid cell: LST1, FCER1G, TYROBP, CTSS, LILRB1, S100A8, and S100A9; and
- platelet: PPBP, PF4, GNG11, and NRGN.

To account for differences in expression distributions across samples, lineage-specific thresholds were estimated within each sample using the zero-of-six reference cells. The primary high-non-T outcome was defined as a B-cell, myeloid, or platelet lineage score exceeding the sample-specific 95th percentile of the corresponding reference distribution, irrespective of the T-cell score. This outcome was additionally partitioned according to T-cell signal. Retained-T/high-non-T cells had T-cell scores at or above the sample-specific reference 25th percentile, whereas low-T/high-non-T cells had T-cell scores below this threshold. More stringent descriptive definitions required T-cell scores above the sample-specific 75th or 95th percentile. These expression patterns were interpreted as heterotypic-like evidence rather than proof of a physical heterotypic doublet.

Natural killer markers were excluded from the strict non-T definition because cytotoxic T cells can physiologically express genes such as NKG7, GNLY, KLRD1, PRF1, and CTSW. An expanded definition including natural killer signal was reported separately.

### 2.9 In silico heterotypic-doublet controls

To assess whether transcript dilution could explain why heterotypic doublets retain moderate rather than extreme T-cell scores, we generated within-sample in silico T–B, T–myeloid, and T–T pairs. Source cells were drawn with replacement and assigned to lineages using Azimuth level-1 annotations. T-cell donors were TCR-positive, non-flagged zero-of-six cells; B-cell and myeloid donors were TCR-negative zero-of-six cells, thereby avoiding non-T donors with discordant TCR evidence. For each eligible sample and pair type, 200 pairs were generated; self-pairing was prohibited for T–T pairs. Raw marker-gene counts and total UMI counts from the two source cells were summed, after which lineage scores were recalculated using the same normalization, marker sets, and sample-specific thresholds as in the observed cell analysis. We summarized high partner-lineage signal irrespective of the T-cell score, low T-cell signal plus high partner signal, and retained T-cell signal plus high partner signal. Definitions using the 75th and 95th T-cell percentiles were retained as sensitivity analyses. The simulation was intended to examine how the scoring rule behaves for known mixtures, not as a generative model of the complete experimental process.

### 2.10 Cell-type annotation

Azimuth level-1 annotations (Hao et al., 2021) were used to compare the proportion of cells assigned to non-T-cell lineages between the unanimous computational-doublet group and the zero-of-six reference group. Because cytotoxic T cells share natural-killer-associated programs, non-T annotations were summarized both excluding and including NK labels.

These annotations were generated before computational doublets were removed and were therefore used as descriptive supporting evidence rather than as an independent ground truth.

### 2.11 Statistical analysis

Continuous lineage scores were compared between the target and reference cells using two-sided Mann–Whitney *U* tests. Differences in the proportions of cells exceeding the sample-specific lineage thresholds were initially assessed using Fisher’s exact tests. Benjamini–Hochberg correction was applied within each family of multiple comparisons.

To control for sample composition, associations between unanimous computational-doublet status and high lineage-marker signals were estimated using Cochran–Mantel–Haenszel analyses stratified by sample. Common odds ratios, 95% confidence intervals, and corresponding *P* values were reported. Participant-stratified analyses were performed as sensitivity analyses. Homogeneity of the stratum-specific odds ratios was assessed using the Breslow–Day test.

Because cell-level stratified analyses do not fully eliminate within-sample correlation, uncertainty for the principal prevalence and TCR-evidence contrasts was additionally quantified using 2,000-replicate cluster bootstrap. Samples and participants were resampled as the independent clustering units in separate analyses, and percentile 95% confidence intervals were reported.

Robustness of the stratified odds ratios to individual samples was assessed using leave-one-sample-out analysis, in which the sample-stratified model was repeatedly refitted after excluding one sample at a time. Statistical tests were two-sided, and adjusted *P <* 0.05 was considered statistically significant.

The six doublet-detection pipelines were executed using R 4.3.2 and Python 3.10.20. The new lineage, chain-level, and robustness analyses were executed in a separate Python environment using anndata 0.13.4, Scanpy 1.12.5 (Wolf et al., 2018), pandas 3.1.0, NumPy 2.5.4, SciPy 1.18.1, statsmodels 0.15.0, scikit-learn 1.9.1, and Matplotlib 3.11.2.

## 3 Results

### 3.1 Study cohort and cross-modal evaluation framework

The analysis comprised 96,003 cells from 44 samples and 29 participants, including 70,959 cells with detectable TCR information. Among TCR-detected cells, 6,208 carried the pipeline-derived TCR potential-multiplet flag. The overall design is summarized in Fig. 2.

**Figure 2:**
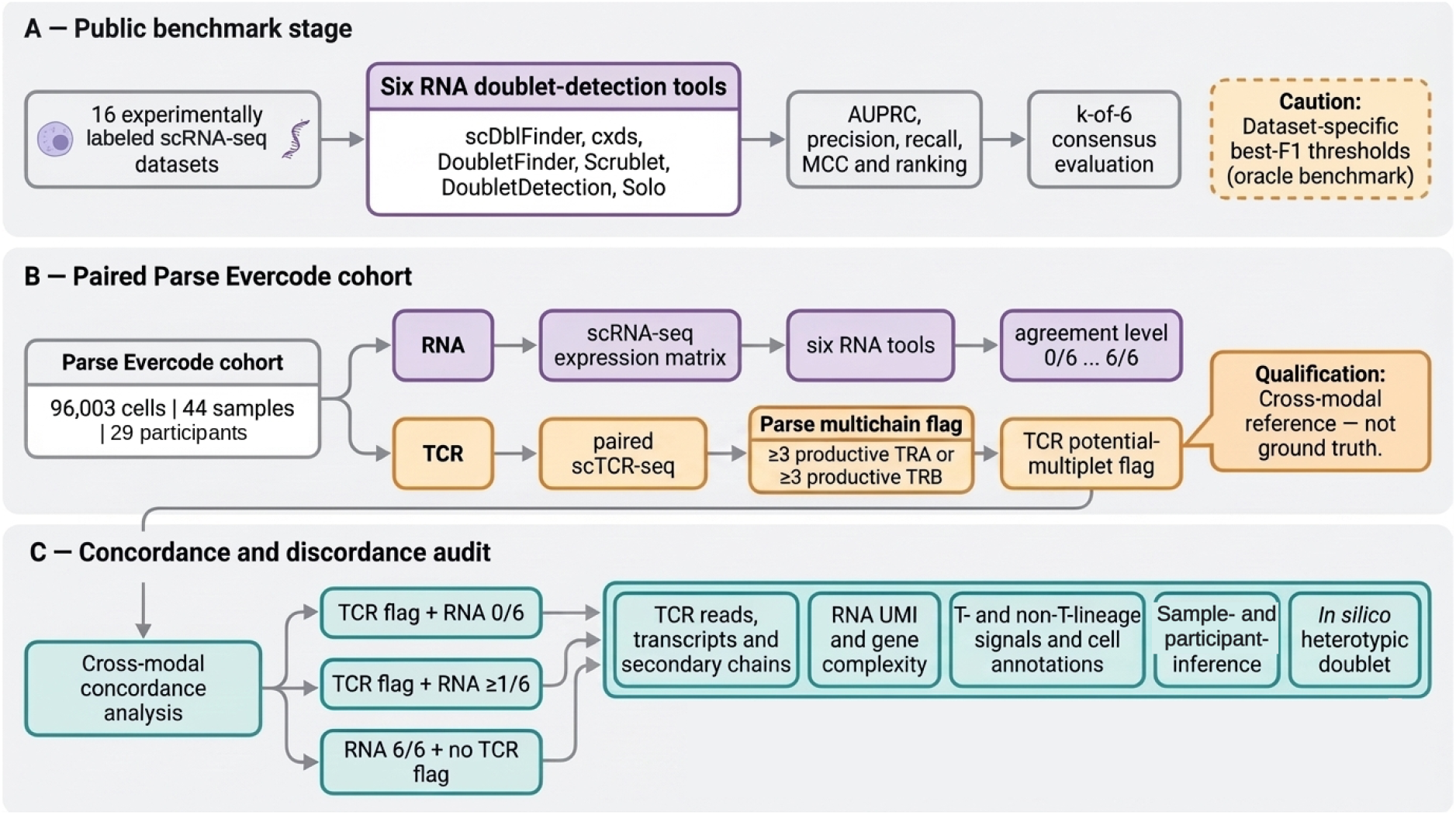
Study design and cross-modal evaluation framework. (A) Six RNA-based doublet-detection tools were evaluated in 16 experimentally labeled public datasets; binary calls in this benchmark used dataset-specific best-F1 thresholds and therefore represent an oracle benchmark. (B) In the paired Parse Evercode cohort, RNA-tool agreement was compared with a Parse-derived TCR potential-multiplet flag based on multichain evidence. The TCR flag was treated as a cross-modal reference, not ground truth. (C) Concordant and discordant groups were audited using chain-level TCR evidence, RNA complexity, lineage signals, cell annotations, sample- and participant-aware inference, and in silico heterotypic-doublet controls.

### 3.2 TCR flags rose with RNA-tool agreement

We first asked whether the TCR flag, which is independent of the features used by RNA-based algorithms, became more frequent as more RNA-based methods called a cell a doublet. Among cells with detectable TCR information, flag prevalence increased from 2.58% (1,453/56,367; Wilson 95% CI, 2.45–2.71%) at zero RNA calls to 7.04% at one, 27.29% at two, 38.62% at three, and a peak of 50.21% (1,561/3,109; 48.45–51.97%) at four calls (Fig. 3A). Flag prevalence was therefore approximately 19-fold higher among cells called by four methods than among cells called by none, providing support for RNA-based consensus from a modality that these algorithms do not use. Pooling all cells called by four or more methods gave a similar estimate (48.34%; 2,273/4,702; Wilson 95% CI, 46.91–49.77%; 18.8-fold), so this contrast does not depend on selecting the peak agreement level.

**Figure 3:**
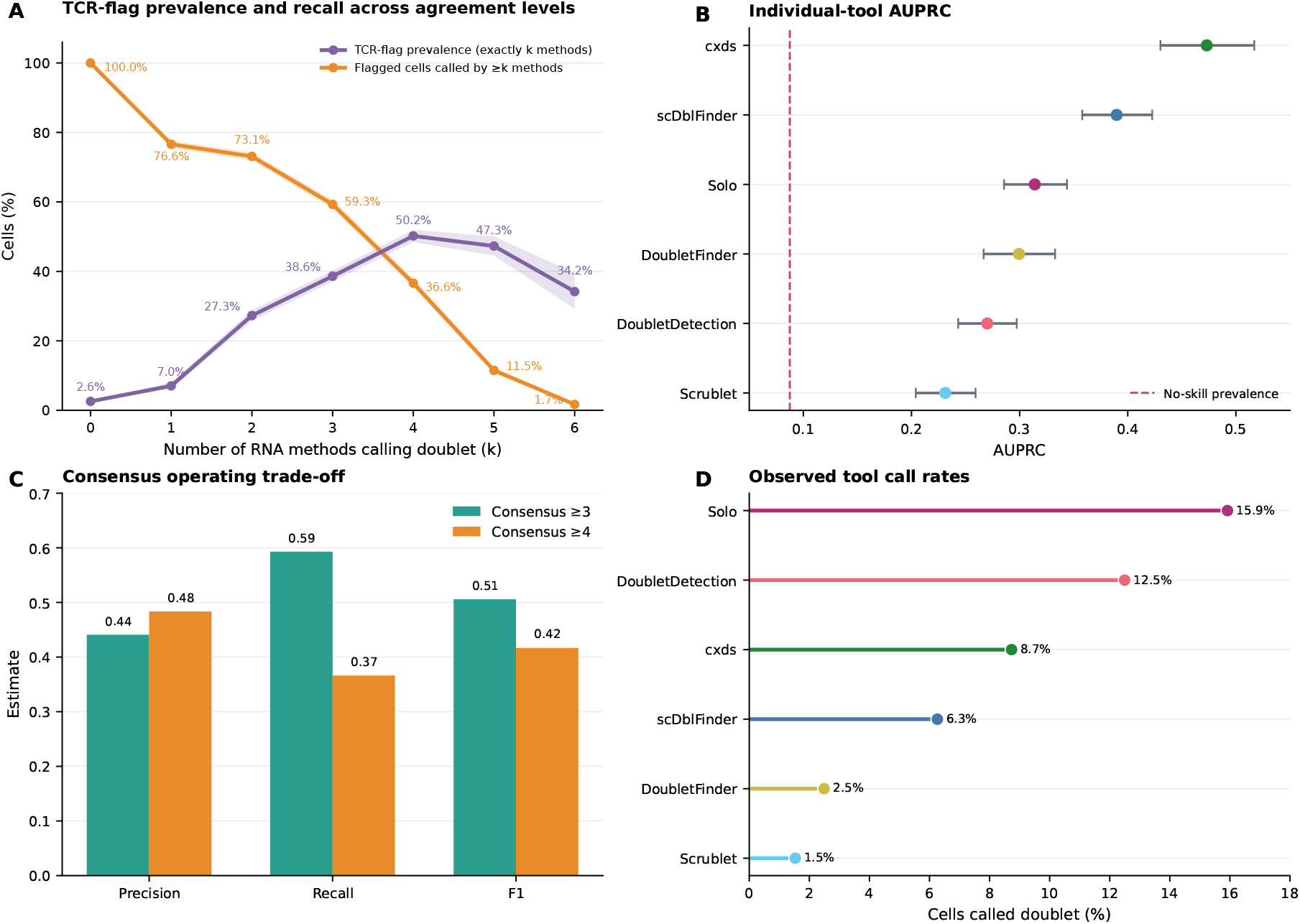
RNA-tool agreement and operating characteristics relative to the TCR potential-multiplet flag. (A) Prevalence of the TCR flag across exact RNA-tool agreement levels (purple) and the percentage of all 6,208 flagged cells called by at least *k* methods (orange); shaded bands are Wilson 95% confidence intervals. (B) Mean sample-level AUPRC for the six tools, with 2,000-replicate sample-cluster bootstrap confidence intervals. (C) Precision, recall, and F1 for consensus thresholds of at least three and at least four positive tools. (D) Observed tool-specific call rates in the Parse cohort. The TCR flag is a cross-modal reference rather than complete ground truth.

Because TCR transcripts are captured from the same mRNA pool, barcodes with high RNA content could be both more likely to be called by RNA-based methods and more likely to yield additional TCR chains. Flag prevalence among cells called by no method varied little across within-sample UMI quintiles (2.1% in the lowest and 3.2% in the highest), but 96% of cells called by four or more methods fell in the highest quintile, so the agreement gradient could not be examined at low RNA depth (Fig. 4). Within the highest quintile, flag prevalence was 15.6 times higher among cells called by four or more methods than among cells called by none (5.2 times in the fourth quintile). In logistic models with sample fixed effects, adjustment for within-sample UMI quintile attenuated the odds ratio per additional positive method from 2.29 (sample-cluster bootstrap 95% CI, 2.13–2.44) to 1.71 (1.61–1.82); adjustment for detected-gene quintile gave 1.70 (1.60–1.81). Part of the cross-modal agreement may therefore reflect shared dependence on RNA content, but a substantial association remained after adjustment. Because multiplets themselves carry more RNA than single cells, RNA depth is partly a consequence of multiplet status rather than only a confounder, so the adjusted estimate may understate the association. Prevalence then declined to 47.30% at five calls and 34.18% (108/316; 29.17–39.57%) at six, indicating that RNA-tool agreement and TCR-defined multiplet status are not equivalent. Part of this decline is consistent with the T–non-T-like profiles among unanimous calls described in Section 3.4, which a chain-count rule cannot identify.

**Figure 4:**
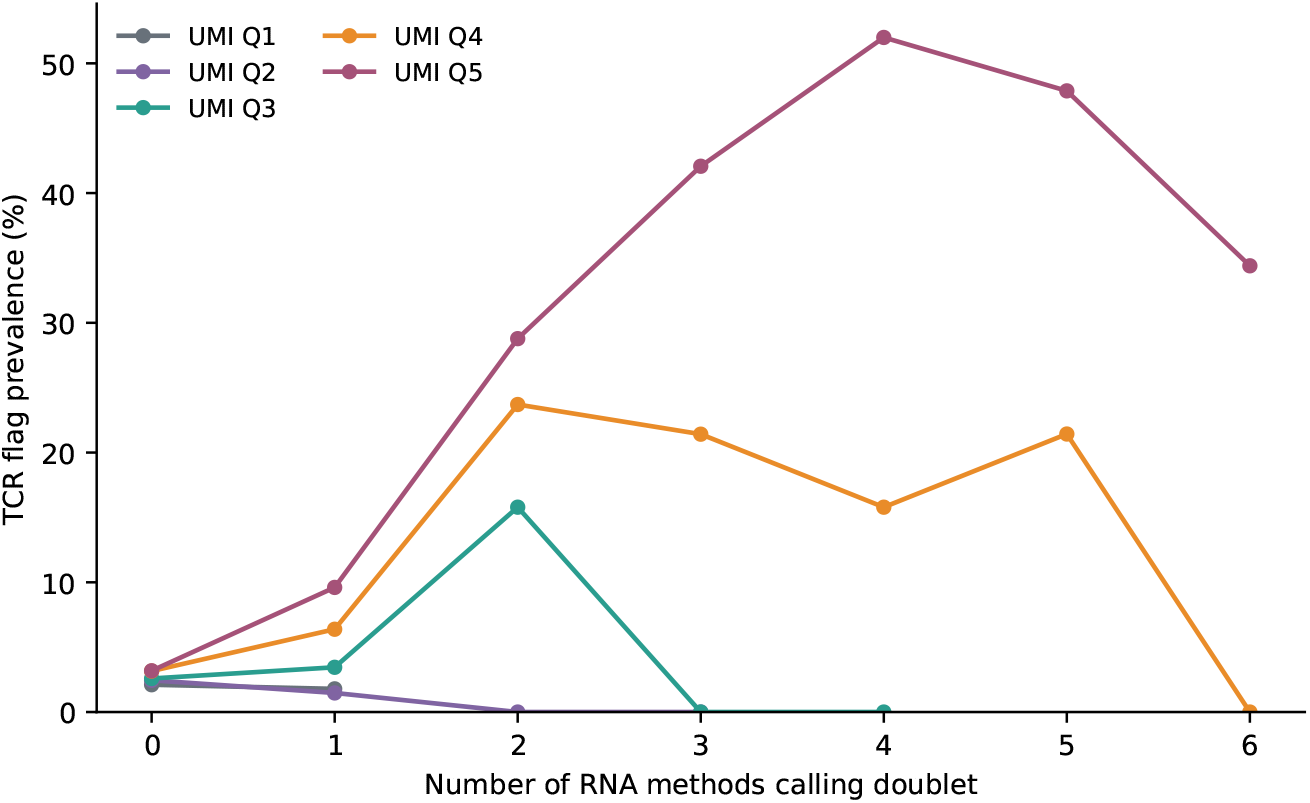
TCR potential-multiplet prevalence by RNA-tool agreement level within quintiles of within-sample total-UMI percentile rank, among 70,959 TCR-informative cells. Agreement levels with no cells in a quintile are omitted; cells called by four or more methods occurred almost exclusively in the highest quintile.

Requiring more methods to agree also came at a cost. Of the 6,208 flagged cells, 76.6% were called by at least one method, 59.3% by at least three, and 36.6% by at least four (Fig. 3A). Requiring four rather than three positive methods increased precision against the TCR flag by 4.25 percentage points (paired sample-cluster bootstrap 95% CI, 2.15–6.38) but reduced recall by 22.68 points (17.59–26.92) and F1 by 8.91 points (5.59–11.65) (Fig. 3C); participant-cluster intervals were similar. In the public benchmark, consensus *≥* 4 achieved the highest mean MCC (0.533). Because that benchmark used label-optimized thresholds, we use *≥* 4 as a reference operating threshold rather than a universally optimal choice; the appropriate threshold may depend on the dataset and on the relative costs of false-positive and false-negative calls.

Individual methods also differed substantially. Call rates ranged from 1.53% for Scrublet to 15.92% for Solo, so differences in recall partly reflected different call budgets rather than sensitivity alone. cxds achieved the highest mean sample-level AUPRC (0.473; sample-cluster bootstrap 95% CI, 0.430–0.517), DoubletFinder the highest precision (0.469), and Solo the highest recall (0.718); no method performed best on every metric (Fig. 3B,D). Tool rankings by mean AUPRC against the TCR flag showed no evidence of concordance with rankings by mean AUPRC across the public benchmark datasets (Spearman’s *ρ* = *−*0.086, *P* = 0.872; six tools), an exploratory result consistent with reference-dependent performance.

### 3.3 TCR-only flags lacked transcriptomic evidence of multiplet content

We next examined discordance in the first direction: the 1,453 of 6,208 TCR potential multiplets (23.4%) that received no positive RNA call. Conventional multiplets would be expected to carry roughly twice the RNA content of a single cell; these cells did not. Their RNA complexity was close to that of all non-flagged TCR-positive cells (median total UMI, 6,549 versus 6,542; Mann–Whitney *P* = 0.113; median detected genes, 2,579 versus 2,569; *P* = 0.222) and slightly but significantly higher than that of the zero-call reference cells used for the unanimous-call comparisons (median total UMI, 6,549 versus 6,227, about 5% higher; *P* = 1.9 *×* 10*^−^*^16^; median detected genes, 2,579 versus 2,494; *P* = 2.7 *×* 10*^−^*^18^), far short of the twofold excess expected of a conventional multiplet, and their full UMI distribution overlapped that of reference cells (Fig. 5C). Because this group was defined by the absence of RNA calls, and RNA-based methods preferentially call barcodes with high RNA content, this observation is partly a consequence of selection and does not by itself exclude physical multiplets.

**Figure 5:**
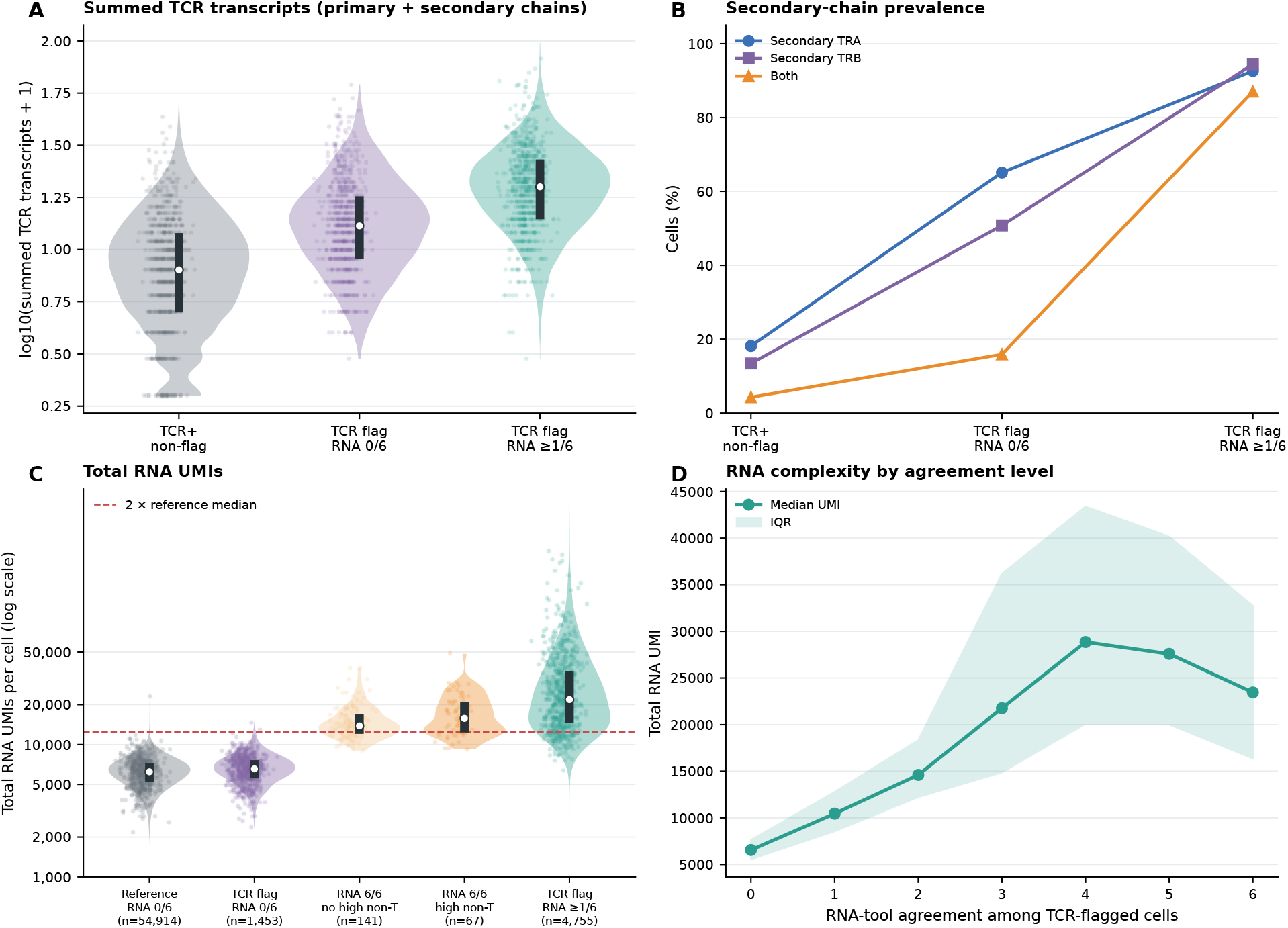
Chain-level and RNA-complexity profiles of TCR audit groups and unanimous RNA calls. (A) Distributions of UMI-deduplicated TCR transcripts summed over the reported primary and secondary chains among non-flagged TCR-positive cells, TCR-flagged cells receiving zero RNA calls, and TCR-flagged cells receiving at least one RNA call. (B) Prevalence of secondary-chain evidence across these groups. Because the flag is defined from chain counts, (A) and (B) describe chain configuration rather than independent validation of the flag. (C) Distributions of total RNA UMIs, on a log scale, among reference cells (TCR-positive, non-flagged, called by no RNA method), TCR-flagged cells receiving zero RNA calls, unanimous RNA calls without the flag split by high non-T-lineage signal, and TCR-flagged cells receiving at least one RNA call; points show all cells in groups of 208 or fewer and a random subset otherwise, white dots medians, and bars interquartile ranges. The dashed line marks twice the reference median. (D) Median total RNA UMI count (with interquartile range) across RNA-tool agreement levels among TCR-flagged cells.

On the TCR side, these cells had more summed TCR transcripts from the reported primary and secondary chains (median, 12 versus 7; difference, 5; sample-cluster 95% CI, 4–6) and more frequent secondary TRB chains (50.8% versus 13.4%; difference, 37.3 percentage points; 34.5–39.9) (Fig. 5A,B); participant-cluster bootstrap intervals led to the same conclusions. Because the flag itself is defined from chain counts, these summaries describe how the cells were flagged rather than independently confirming that they contain more than one cell. Among non-flagged TCR-positive cells, TCR evidence was only weakly correlated with RNA content (median within-sample Spearman’s *ρ*, 0.138 for reads and 0.189 for transcripts), so these comparisons are reported without adjustment for RNA complexity.

By contrast, the 4,755 flagged cells called by at least one RNA method were elevated in both modalities: their RNA complexity was markedly higher (median UMI, 21,856; median detected genes, 5,330), and their TCR signals were also higher (median summed TCR transcripts, 19; difference from the zero-call group, 7; sample-cluster 95% CI, 5–8; secondary TRB, 94.4%). Among flagged cells, median RNA complexity rose from 6,549 UMIs at zero RNA calls to 28,853 at four calls and declined modestly at five and six (Fig. 5D). RNA-tool agreement was also associated with RNA complexity among non-flagged cells, but the association was stronger among flagged cells (within-sample Spearman’s *ρ* for total UMI, 0.784 versus 0.546; difference, 0.238; sample-cluster 95% CI, 0.157–0.307). These concordant cells form the clearest overlap between the two modalities.

### 3.4 RNA-based methods identified T–non-T-like events that lacked a TCR flag

We then examined discordance in the opposite direction. If unanimous RNA calls that lacked a TCR flag included mixtures of a T cell with a non-T cell, they should carry transcripts of the non-T partner. We therefore compared the 208 TCR-detected, non-flagged cells called by all six RNA methods with 54,914 TCR-detected, non-flagged cells called by none (reference cells). Each cell was scored for B-cell, myeloid, and platelet marker expression and classified as having high non-T signal if any of these scores exceeded the 95th percentile of reference cells in the same sample (Methods).

Sixty-seven of the 208 unanimous calls (32.2%) had high non-T signal, compared with 10.5% of reference cells. The reference value is a baseline set by the thresholds themselves: by construction, about 5% of reference cells exceed each of the B-cell and myeloid thresholds. High non-T signal was therefore about three times as prevalent among unanimous calls (32.2% versus 10.5%; sample-stratified common odds ratio, 4.08; 95% CI, 3.03–5.50). The association did not differ detectably across samples (Breslow–Day *P* = 0.15) and was similar when stratified by participant (OR, 4.11; 3.06–5.52), when each sample was omitted in turn (OR range, 3.61–4.46), when B-cell, myeloid, and platelet signals were analyzed separately (OR, 5.26, 3.15, and 11.07, respectively), and in an unstratified Fisher’s exact test (OR, 4.06). A complementary reference-based annotation agreed: 4.8% of unanimous calls versus 0.6% of reference cells received a non-T, non-NK Azimuth label (B cell, monocyte, dendritic cell, or other), and all ten such unanimous calls were labeled B cells (sample-stratified OR, 7.40; 3.74–14.64; participant-stratified OR, 8.73; 4.48–16.98; Fig. 6A). NK labels were excluded from this comparison because cytotoxic T cells share NK-associated programs; including them gave 15.9% versus 2.1% (OR, 8.27; 5.56–12.30).

**Figure 6:**
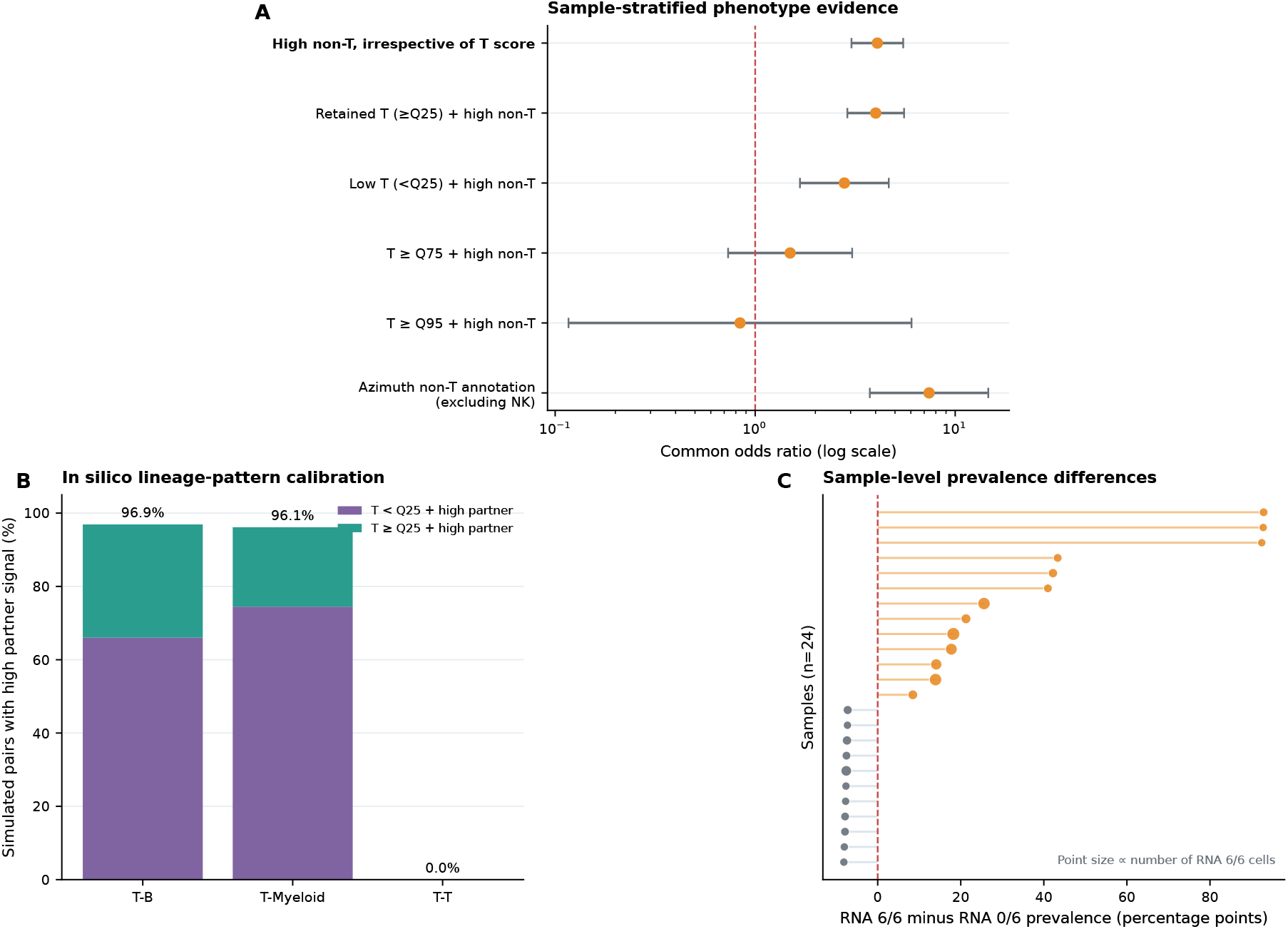
Phenotypic audit of unanimous RNA doublet calls lacking the TCR potential-multiplet flag. (A) Sample-stratified phenotype associations, reported as common odds ratios with 95% confidence intervals; the first row is the primary outcome. (B) In silico T–B, T–myeloid, and T–T controls. Bar height is the percentage of simulated pairs with high partner-lineage signal, divided into pairs with T-cell scores below (low T) or at or above (retained T) the sample-specific reference 25th percentile. (C) Sample-level differences in the prevalence of high non-T-lineage signal with retained T-cell signal, for the 24 samples containing unanimous calls. In (A), the Azimuth non-T category excludes NK labels; estimates including NK labels are reported in the text. Evidence supports a heterotypic-like subset but does not identify the physical composition of individual cells.

The enrichment was present whether the T-cell score of a unanimous call remained typical or fell below the reference 25th percentile, but there was no clear evidence of enrichment when a high T-cell score was also required (Fig. 6A), a pattern consistent with the in silico mixtures described below. For the retained-T/high-non-T definition, the within-sample difference was positive in 13 of the 24 samples that contained unanimous calls; in the remaining 11, none of the unanimous calls met that definition (Fig. 6C).

To examine how the lineage scoring rule behaves for known mixtures, we summed the counts of T cells with those of TCR-negative B cells or myeloid cells from the same sample. High partner-lineage signal was recovered in 96.9% of in silico T–B and 96.1% of T–myeloid pairs, but most pairs (66.0% and 74.5%) fell below the reference 25th percentile of the T-cell score, and almost none exceeded the 75th percentile. Combining two cells therefore dilutes the T-cell signal, which is why high non-T-lineage signal, rather than simultaneously high T-cell and non-T-cell scores, was used as the primary phenotype (Fig. 6B). These simulations describe the behavior of the scoring rule and do not establish that individual observed cells were physical heterotypic doublets.

The remaining two-thirds of unanimous calls (141 cells) lacked high non-T signal but carried about twice the RNA content of reference cells (median total UMI count, 13,856 versus 6,227; ratio, 2.2; sample-cluster bootstrap 95% CI, 2.12–2.42), ranking at a median within-sample percentile of 96% among TCR-positive, non-flagged cells. The 67 unanimous calls with high non-T signal also carried elevated RNA content (median total UMI count, 15,773; ratio, 2.5; sample-cluster bootstrap 95% CI, 2.12–2.99; median within-sample percentile, 97%), so both subsets of unanimous calls, unlike flag-only barcodes, carried roughly twice the RNA content of reference cells (Fig. 5C).

### 3.5 TCR and RNA evidence captured complementary event classes

Taken together, three event classes emerged. TCR-flagged cells not called by any RNA method had single-cell-like RNA content but more TCR chains; flagged cells called by RNA methods were elevated in both RNA content and TCR signal; and unanimous RNA calls without the flag were enriched for non-T-lineage signal or carried about twice the RNA content of reference cells. RNA content therefore separated the two directions of discordance: barcodes flagged only by TCR carried single-cell-like RNA content, whereas unanimous RNA calls without the flag carried roughly twice as much (Fig. 5C). Each modality contributed events that the other did not, and the two forms of evidence are complementary rather than interchangeable.

## 4 Discussion

In this comparison, in which RNA and TCR were profiled from the same Parse Evercode barcodes, paired scTCR-seq added information that RNA-based doublet detection alone could not provide. Flag prevalence rose roughly 19-fold from cells called by no RNA method to cells called by four, and 4,755 of the 6,208 flagged cells (76.6%) were called by at least one method, providing support for RNA-based consensus from a modality that these algorithms do not use. This support is not entirely independent of capture depth: adjustment for within-sample RNA depth reduced the odds ratio per additional positive method from 2.29 to 1.71, and the agreement gradient could be assessed only among barcodes with high RNA content. The two modalities nonetheless disagreed in structured ways, and each direction of disagreement exposed a limitation of the three-chain rule.

In the first direction, the 1,453 flagged cells that no RNA method called had RNA content similar to that of single T cells, yet they carried more TCR transcripts and more frequent secondary TRB chains. Because the flag is defined from chain counts, these chain-level differences describe how these cells were flagged rather than independently confirming that they contain a second cell. A multiplet of two typical T cells would be expected to carry roughly twice the RNA content of a single cell, and this group did not show such an excess. Because the group was selected for the absence of RNA calls, this does not exclude physical multiplets, but it indicates that their multichain signal is not accompanied by a conventional multiplet transcriptome. Dual-receptor T cells are also insufficient on their own: because each TCR locus has two alleles, a single T cell is expected to express at most two productive chains of each type, below the three-chain threshold. The remaining explanations are TCR-specific technical events, such as ambient or misassigned TCR transcripts or sequence variants of one chain called as distinct chains, and multiplets in which one partner contributes little RNA. In either case, RNA-based methods have little information with which to call them, and they are better regarded as unconfirmed flags than as doublets missed by RNA-based methods. Flagged cells called by at least one RNA method, by contrast, were elevated in both RNA content and TCR signal, and their RNA content rose with RNA-tool agreement; RNA-based methods therefore appear to recognize the subset of multichain events that also carry a transcriptomic abnormality.

In the opposite direction, unanimous RNA calls without the flag were enriched for B-cell, myeloid, or platelet signal and for B-cell Azimuth annotations. Mixtures of a T cell with a non-T cell would be expected to show this pattern and would not be flagged, because they need not contain additional TCR chains. The in silico mixtures showed that combining two cells dilutes the T-cell signal, so a true heterotypic event need not exceed a high T-cell threshold; definitions requiring simultaneously high T-cell and non-T-cell scores were accordingly insensitive. These data do not, however, establish the physical identity of individual cells. Non-T cells carrying ambient or misassigned TCR transcripts, platelet–leukocyte aggregates, and annotation discordance could produce similar profiles, and we therefore describe these cells as heterotypic-like rather than as verified heterotypic doublets. Only about one-third of unanimous calls showed high non-T-lineage signal. The remainder carried roughly twice the RNA content of reference cells, which is compatible with T–T multiplets below the three-chain threshold, such as two conventional T cells contributing two TRA and two TRB chains. Because RNA-based methods preferentially call barcodes with high RNA content, however, large single cells cannot be excluded. Secondary-chain evidence fits neither simple model cleanly. A mixture of one T cell with a B cell or myeloid cell should not add TCR chains, yet unanimous calls with high non-T signal carried secondary TRA chains in 37.3% and secondary chains at both loci in 22.4%, compared with 15.5% and 1.7% of reference cells. Conversely, T–T multiplets would be expected to carry secondary chains at both loci frequently, yet only 17.0% of the remaining unanimous calls did. Possible explanations include multiplets containing more than one T cell together with a non-T cell, greater capture of ambient TCR transcripts in barcodes with high RNA content, and incomplete TCR capture in T–T multiplets; the present data cannot distinguish among them.

Consensus calling offers a transparent summary of heterogeneous tool outputs, but the threshold of four positive methods should not be considered universally optimal. The supporting public benchmark used dataset- and method-specific thresholds optimized against experimental labels, whereas the Parse cohort used package-derived or prespecified operating thresholds. Requiring four rather than three methods produced a modest gain in precision against the TCR flag at the cost of a much larger loss in recall, and flag prevalence rose gradually with agreement rather than jumping at a single threshold. The component methods are also correlated, their call rates differ several-fold, and each experimental reference favors particular multiplet subtypes. Consensus count is therefore best retained as graded evidence, together with the identities of the component methods, rather than converted into a single permanent exclusion rule. The lack of concordance between tool rankings based on the TCR flag and on public benchmarks, although based on only six methods, is consistent with this reference dependence, and the TCR-based ranking should not be read as general superiority of any single method.

In practice, these results suggest that, in immune-profiling datasets with paired TCR sequencing, the TCR potential-multiplet flag and RNA-based consensus should be used together. Events supported by both modalities provide the strongest candidates for exclusion. Flag-only events, which lacked RNA evidence of multiplet content, are better treated as candidates for review than as automatic exclusions, whereas unanimous RNA calls should not be retained merely because they lack a flag, since the chain-count rule cannot detect T–non-T mixtures.

This study benefits from paired modalities generated in the same 44 samples, per-sample execution of six tools, an external benchmark, chain-level summaries, and sample- and participant-cluster bootstrap intervals for the principal prevalence, operating-characteristic, and chain-level contrasts. Several limitations remain. The pipeline-derived flag is not complete ground truth, is restricted to TCR-detected cells, and may include technical events such as ambient TCR transcripts or chain sequence variants; conversely, a multiplet of two conventional T cells, with two TRA and two TRB chains, does not satisfy a three-chain rule. Only summarized primary and secondary chain fields were available, so the additional chains that triggered the flag could not be inspected directly, and independent experimental validation, such as cell hashing or genotype demultiplexing, was not available. The six RNA methods are not independent votes, and DoubletDetection did not meet its convergence criterion at 50 iterations in 11 of 44 samples. The marker-panel analysis cannot establish the physical identities of contributing cells, and its sample-specific thresholds follow common conventions rather than an external standard. Finally, the PBMC cohort, which was enriched for Graves’ disease and antithyroid drug-induced agranulocytosis, and the Parse Evercode platform may limit generalization to other tissues and technologies.

Two extensions would address these gaps. Inspecting the complete chain set would test whether the additional chain in flag-only barcodes is a near-copy of another chain, as expected from a sequencing error, or a distinct receptor. More generally, when no reference is complete, latent class models that treat each RNA-based method and the TCR flag as imperfect measurements of a shared latent multiplet status (Hui and Walter, 1980) could combine these sources formally, provided that the dependence among RNA-based methods is modeled explicitly.

In conclusion, paired scTCR-seq is a valuable complementary reference for doublet detection. TCR multichain evidence supported RNA-based consensus, but flags without RNA support carried single-cell-like RNA content and are best reviewed rather than removed, whereas RNA-based methods identified T–non-T-like mixtures and high-RNA events that the TCR flag cannot capture. In immune-profiling studies with paired TCR data, combining both modalities and retaining graded evidence from each offers a practical path toward more complete doublet detection.

## Data availability

The single-cell sequencing data were generated in a previously described cohort (Huang et al., 2025) and are not publicly available because participant consent did not cover deposition in public repositories. Data access is described in Huang et al. (2025).

## Code availability

Analysis code used to execute and compare the six RNA-based doublet-detection methods, construct multi-tool consensus calls, perform cross-modal comparisons, characterize discordant cells, and generate the reported figures and summary statistics is available from the corresponding authors upon reasonable request.

## Ethics approval

The study involved human participants and was conducted in accordance with the Declaration of Helsinki. The Research Ethics Committee of National Taiwan University Hospital gave ethical approval for this work (Approval No. 201611020RIND). Written informed consent was obtained from all participants.

## Author contributions

Y.-T.W. and S.-H.L. contributed equally to this work. P.-L.C. and C.-Y.C. initiated the study. Conceptualization: P.-L.C., C.-Y.C., Y.-T.W., S.-H.L. Methodology: Y.-T.W., S.-H.L. Software: Y.-T.W., S.-H.L. Formal analysis: Y.-T.W., S.-H.L. Validation: S.-H.L. Visualization: S.-H.L. Data curation: Y.-T.W., S.-K.L. Investigation: Y.-H.H., S.-K.L., I-H.C. Resources: Y.-H.H., S.-K.L., I-H.C., P.-L.C. Supervision: P.-L.C., C.-Y.C. Project administration: C.-Y.C. Funding acquisition: P.-L.C., C.-Y.C. Writing – original draft: S.-H.L. Writing – review & editing: all authors.

## Acknowledgements

We thank all participants who contributed samples to this study, and all clinical and laboratory staff who supported sample collection and processing.

